# Distinct changes in brain and peripheral oxygen levels induced by synthetic opioids of the nitazene family

**DOI:** 10.64898/2026.11.12.766190

**Authors:** Ali Arce, Nicholas Nguyen, Juan L. Gomez, Agnieszka Sulima, Kenner C. Rice, Michael Michaelides, Eugene A. Kiyatkin

## Abstract

Nitazenes are a category of synthetic opioids, some of which have high analgesic potential but also high toxicity, which has historically prevented their development as therapeutic drugs. Like fentanyl, nitazenes induce respiratory depression, a primary factor in their toxicity. Recently, a nitazene derivative called N-desethyl-fluornitrazene (DFNZ) was shown to have high analgesic properties with weak hypoxic effects. To better characterize the hypoxic effects of DFNZ relative to its parent, fluornitrazene (FNZ) and the prototypical nitazene etonitazene, we used amperometric oxygen sensors in freely moving rats to examine oxygen dynamics in the nucleus accumbens (NAc) and the subcutaneous space, using fentanyl as a comparison. All drugs induced similar patterns of subcutaneous hypoxia, with rapid dose-dependent oxygen decreases. In the NAc, all opioids, except DFNZ, at low iv doses (1-5 μg/kg) induced moderate but significant brain oxygen increases. At higher doses (>10 μg/kg), oxygen levels for all opioids, except DFNZ, showed a biphasic response; robust hypoxia followed by a weaker rebound-like hyperoxia. Etonitazene at 30 μg/kg had the largest and most prolonged brain hypoxia compared to the other drugs, without the rebound-like hyperoxia. These findings indicate that etonitazene and FNZ mimic the effects of fentanyl, inducing dose-dependent brain and peripheral hypoxia. In contrast, DFNZ induces very weak hypoxia seen minimally at very high iv doses (300-1000 μg/kg). Our findings substantiate the low hypoxic potency of DFNZ and indicate that not all nitazenes are characterized by high potential for toxicity. As such, the development of nitazenes as analgesics should not be categorically dismissed.

**Highlights:**

- All nitazene drugs tested induced dose-dependent brain and peripheral hypoxia
- Brain hypoxia induced by etonitazene and FNZ was strong and most prolonged
- DFNZ induced very weak brain hypoxia seen minimally at a 300 μg/kg iv dose

## 1. Introduction

Nitazenes, also known as 2-benzyl benzimidazole opioids, are a category of potent synthetic opioids first synthesized in the 1950s, with numerous additional derivatives of this family synthesized over the past 70 years. While highly potent in analgesic tests, the first nitazenes were also found to be highly toxic (Baumann, 2026; Vandeputte et al., 2025; Dow et al., 2026; Ujváry et al., 2021; Skolnick 2022). Therefore, clinical and preclinical efforts to develop nitazenes for analgesia were largely abandoned.

In recent years, many different nitazenes have appeared in recreational drug markets, often as adulterants with fentanyl and other drugs of abuse. Due to their extreme potency, nitazenes are associated with an exceptionally high risk of fatal overdose (Shover et al., 2021; Jones et al., 2018; Schwarz et al., 2025; Prekupec et al., 2017) with respiratory depression as the most dangerous side-effect (Malcolm et al., 2023). Nevertheless, as a distinct synthetic opioid class, nitazenes have not been systematically evaluated in the scientific literature and in particular, the effects of individual nitazene analogs on respiratory depression are unclear.

Most information about drug-induced brain hypoxia relies on the use of plethysmography data, which measures changes in breathing activity, or pulse oximetry, which measures *the saturation of oxygen carried by red blood cells*. In our studies, we have used electrochemical oxygen detection, which can directly assess drug-induced changes in oxygen levels in brain and peripheral tissues (Kiyatkin, 2019). This electrochemical technology was previously used to examine the hypoxic effects of several widely used opioids, including heroin (Thomas et al., 2021), fentanyl (FEN) (Choi et al., 2024, Curay et al., 2023) and oxycodone (Solis et al., 2018). In these prior studies, all tested opioids induced brain hypoxia, but their potency varied greatly depending on the drug, dose, and route of administration.

Recently, it was shown that N-desethyl-fluornitrazene (DFNZ), a novel fluorinated nitazene metabolite has superagonist efficacy, high potency, strong analgesic properties, but surprisingly, relatively weak brain hypoxic effects and low toxicity (Gomez et al., 2026). This dissociation between analgesic efficacy and hypoxia is unusual for an opioid, especially potent and high efficacy agonists, which typically show parallel changes in analgesic and hypoxic effects. The discovery of DFNZ prompted us to systematically examine its hypoxic effects and compare them with other potent nitazenes, including the prototypical nitazene etonitazene (ETO), DFNZ’s parent fluornitrazene (FNZ), and FEN, the most prevalent synthetic opioid drug. As such, here we used the electrochemical technique to examine changes in oxygen responses in the brain (nucleus accumbens or NAc) and peripheral tissue (subcutaneous space) induced by intravenous (iv) bolus injection of ETO, FNZ, DFNZ, and FEN in awake, freely moving rats.

## 2. Materials and Methods

### 2.1 Subjects

28 adult male Long–Evans rats (Charles River Laboratories) weighing 450–520 g at the time of surgery were used in this study. Rats were individually housed in a climate-controlled animal colony maintained on a 12:12 light-dark cycle (07:00 AM light on) with food and water available ad libitum. All procedures were approved by the National Institute on Drug Abuse-Intramural Research Program (NIDA-IRP) Animal Care and Use Committee and complied with the Guide for the Care and Use of Laboratory Animals (NIH, Publication 865-23) and ARRIVE guidelines. Maximal care was taken to minimize the number of experimental animals and any possible discomfort or suffering at all stages of the study.

### 2.2. Overview of the Study

Electrochemical experiments were conducted in 28 freely moving rats, using a total of 102 recording sessions. The protocols of each experiment were identical in all rats. To reveal the role of one factor (drug), all other factors that may potentially affect the results (animal age, weight, and biological sex, procedures of stimulus presentation, the drug dose) were identical in all rats.

### 2.3. Surgical preparations

Surgical procedures for the electrochemical assessment of oxygen have been described in detail in prior studies (Choi et al., 2024; Limiac et al., 2025). Under general anesthesia (ketamine HCl and xylazine HCl 80 and 8 mg/kg, respectively, with additional boosting at 30% of initial doses as necessary), rats were chronically implanted with Pt-Ir oxygen sensors (Model 7002-02; Pinnacle Technology, Inc., Lawrence, KS, USA) in two locations. One sensor was implanted in the medial segment of the NAc (AP +1.2 mm, ML ±0.8 mm, and DV +7.2-7.6 mm from the skull surface) according to coordinates from the rat brain atlas (Paxinos and Watson, 1998), while the other sensor was implanted in the subcutaneous space in the frontal area of the rat’s head. This area is densely vascularized, and the sensor implanted in this area does not move during behavioral activation, allowing for artifact-free electrochemical recording. Both sensors were secured with dental acrylic to three stainless steel screws threaded into the skull. During the same surgery, rats were implanted with a chronic jugular catheter, which ran subcutaneously to the head mount. Rats were allowed a minimum of 5 days of post-operative recovery and at least 3 daily habituation sessions (∼6 h each) to the recording environment. Jugular catheters were flushed daily with 0.2 ml heparinized saline to maintain patency.

### 2.4. Electrochemical detection of oxygen

For *in vivo* oxygen detection, we used Pinnacle oxygen sensors coupled with high-speed amperometry. The principles of electrochemical oxygen detection, construction of oxygen sensors, and their calibration have been described in detail in prior studies (Kiyatkin, 2019; Thomas et al., 2021).

### 2.5. Experimental procedures

In this study, we used rats with chronically implanted oxygen sensors in two locations (the NAc and subcutaneous space) and chronic iv catheters for drug delivery. Each rat underwent several recording sessions (up to 6 sessions for a total of 102 total recording sessions across the entire experiment), and their numbers were dependent upon the quality of recording by each sensor and catheter’s patency. There were cases in which one sensor exhibited artifacts, while the other produced artifact free recordings. Therefore, the number of tests for each stimulus differed from the number of rats, in which these tests were performed.

At the onset of each recording session, rats were briefly anesthetized (<2 min) with isoflurane and oxygen sensors were connected to the recording instruments via an electrically shielded flexible cable and a multi-channel electrical swivel. A catheter extension mounted on the cable was used to allow stress- and cue-free drug delivery from outside the cage. After stabilization of baselines, rats were exposed to iv delivery of either fentanyl citrate ((FEN); Hospira Inc., Lake Forest, IL), ETO, FNZ, or DFNZ.

In most cases, rats were tested with a single drug at different ascending doses within a single recording session. Each drug delivery occurred when the rats were in a quiet resting state with no overt movements and when electrochemical currents were relatively stable. Durations of between-injection time intervals were based on previous experience, pilot tests, and were between 40 and 120 min depending on the drug dose, restoration of baseline for electrochemical currents, and observed behavioral effects. When testing was completed, each rat was lightly anesthetized by isoflurane and disconnected from the recording instruments. Rats were allowed to recover from anesthesia, and their jugular catheters were flushed with heparinized saline before being returned to the animal colony. The same protocols were maintained during the repeated daily sessions. In a few sessions, we were unable to maintain this strict protocol due to technical issues with the recording quality of electrochemical sensors and catheter patency.

### 2.6. Histological verification of electrode placements

After experiments were complete, rats were deeply anesthetized with isoflurane, decapitated, and their brains were extracted and stored in 10% formalin solution. Then, the brains were cut and analyzed for verification of the locations of cerebral implants and possible tissue damage at the area of electrochemical recording.

### 2.7. Data analysis

Electrochemical data were analyzed in a slow (1-min) and a rapid (4-s) time resolution. Because each individual sensor differed in substrate sensitivity, currents were first converted into concentrations (μM) based on sensitivity calibrations provided by the manufacturer and then into percent (%) changes in oxygen concentration. Values from one-minute and four-seconds prior to the drug injection were averaged and set as the 100% baseline for slow and rapid time-course analyses, respectively. One-way repeated-measures ANOVA (followed by Dunnett’s multiple comparisons test) was used to evaluate statistical significance of drug-induced oxygen responses. Drug-induced hypoxic responses were quantified by the magnitude of the maximal drop in electrochemical current and by the duration of the signal decrease (until crossing a pre-injection baseline). We also calculated area under the curve (AUC) as an integral parameter of oxygen response. These parameters were determined in the NAc and SC space of each drug from baseline-corrected percent changes, and group means ± SEM were compared using Welch’s unequal variances t-test. For text clarity, quantitative results of statistical evaluations are shown in the Supplemental Materials (Supplementary Tables S1-S5).

## 3. Results

### 3.1. Brain and peripheral oxygen fluctuations induced by fentanyl

Consistent with our previous studies (Thomas et al., 2021; Curay et al., 2023; Choi et al., 2024), iv FEN induced complex changes in oxygen levels, which differed depending on the dose and the recording location (Figure 1A; see Supplementary table S1 for results of statistical evaluations). When injected at the lowest, 1 µg/kg dose, FEN had no effects on oxygen levels in both the NAc and subcutaneous space. At a higher, 3 µg/kg dose, FEN induced minimal oxygen changes with slight *increases* in the NAc and larger but transient decreases in the subcutaneous space. The same pattern of oxygen responses was also seen after FEN injection at a moderate,10 µg/kg dose. However, these changes were more pronounced and prolonged. At a higher, 30 µg/kg dose, FEN induced biphasic oxygen responses in the NAc, with a rapid but transient decrease followed by more prolonged rebound-like increase. In the subcutaneous space the response was clearly monophasic with a rapid and robust decrease (30-40% of baseline) and slow return to the pre-injection baseline. The same pattern of oxygen responses was also seen after FEN injection at the largest, 90 µg/kg dose. At this dose, some rats showed convulsive movements and gasping, preventing further drug testing at larger doses.

**Fig. 1.**
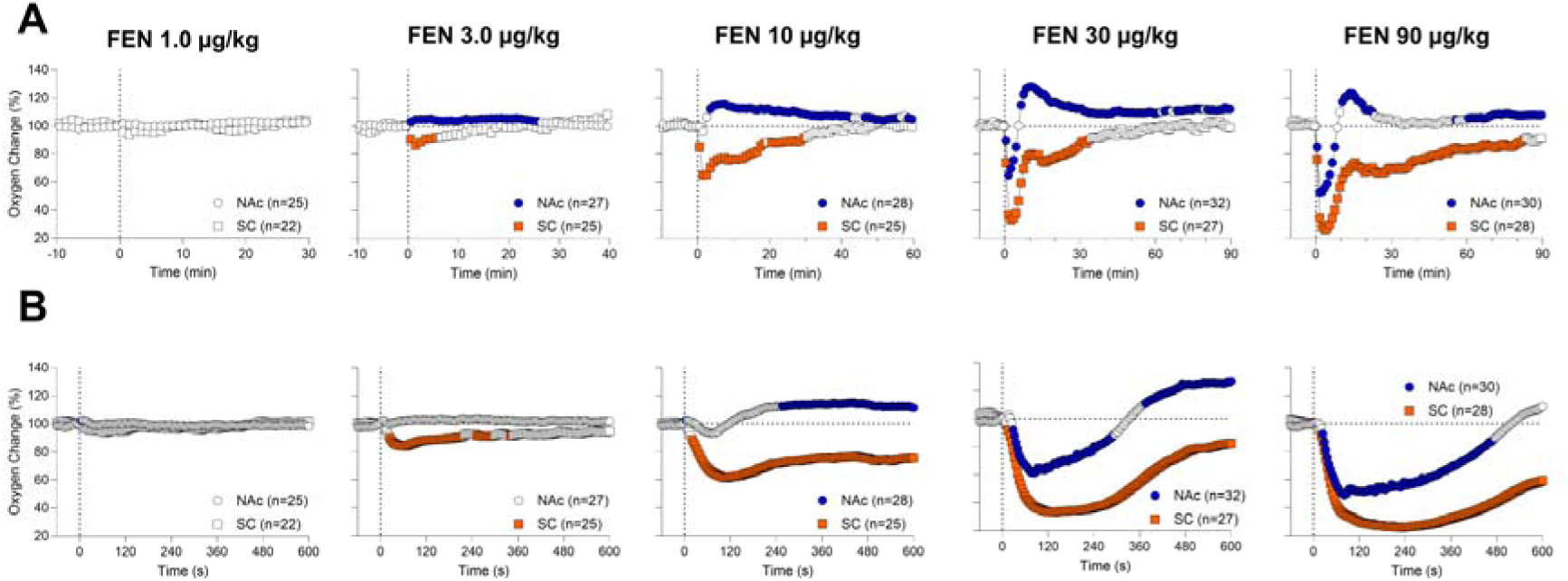
Mean (±SEM) changes in oxygen levels induced by iv fentanyl in the NAc and subcutaneous space at different doses. A shows data analyzed with slow (1-min) time resolution. B shows data analyzed with rapid (4-s) time resolution. Results of statistical evaluations are shown in Supplementary materials (S1). Filled symbols show values significantly different from baseline (p<0.05). n = numbers of averaged tests.

For more precise assessment of FEN-induced oxygen fluctuations, data were analyzed with high, 4-s temporal resolution for the first 10 min after injection onset (Figure 1B). At lower doses, oxygen responses in the subcutaneous space had much shorter onset latencies than in the NAc, but latencies between NAc and subcutaneous space were identical at higher FEN doses. High-resolution analyses also revealed a minor transient fall in oxygen levels in the brain, which was seen within the first two minutes after fentanyl injection at a 10 µg/kg dose. With both higher doses, decreases in the NAc rapidly reached their nadir (∼50 and 60% of baseline at 80-100 s) but their mean durations were relatively small, varying between ∼5-8 min after the injection onset. Compared to the NAc, oxygen decreases induced by large doses of fentanyl in the subcutaneous space were also larger in magnitude (35-25% of baseline at 200-240 s at 30 and 90 µg/kg doses), peaked later, and were maintained for much longer time post-injection.

### 3.2. Brain and peripheral oxygen fluctuations induced by etonitazene (ETO)

The same protocol was used to examine the effects of ETO on central and peripheral oxygen levels, and the effects of this drug were compared to the effects of FEN. Due to its high toxicity (Baumann, 2026; Vandeputte et al., 2023; Dow et al., 2026), the range of tested doses was three-fold lower than FEN. At the largest dose of ETO (30 µg/kg), most rats showed convulsions, extreme pose changes, strong tail erection, and high muscular tone.

ETO induced complex dose-dependent changes in oxygen with profound differences between central and peripheral recording locations (Figure 2A and B; see Supplementary table S2 for results of statistical evaluations). At the lowest 0.33 µg/kg dose, ETO has no effect of oxygen levels in both locations. However, at a 1.0 µg/kg dose, ETO significantly increased oxygen levels in the NAc and decreased them in the subcutaneous space. The magnitude of these responses was within the range of spontaneous oxygen fluctuations (±15% above and below of baseline). Surprisingly, despite evident behavioral sedation, no NAc oxygen decreases were seen at a higher, 3.0 µg/kg dose. In this case, oxygen levels tonically increased and subcutaneous space oxygen levels tonically decreased. At a higher, 10 µg/kg dose, ETO induced very weak decreases in NAc oxygen levels coupled with profound and prolonged decreases in the subcutaneous space. At the largest, 30 µg/kg dose, ETO decreased oxygen in both recording locations. Oxygen decreases in the NAc (mean=66.23% of baseline, SD=7.71%) were smaller than decreases in the subcutaneous space (mean=37.55% of baseline, SD=17.19%). Brain oxygen response induced by ETO at a 30 µg/kg dose was monophasic, with virtually no rebound-like second phase.

**Fig. 2.**
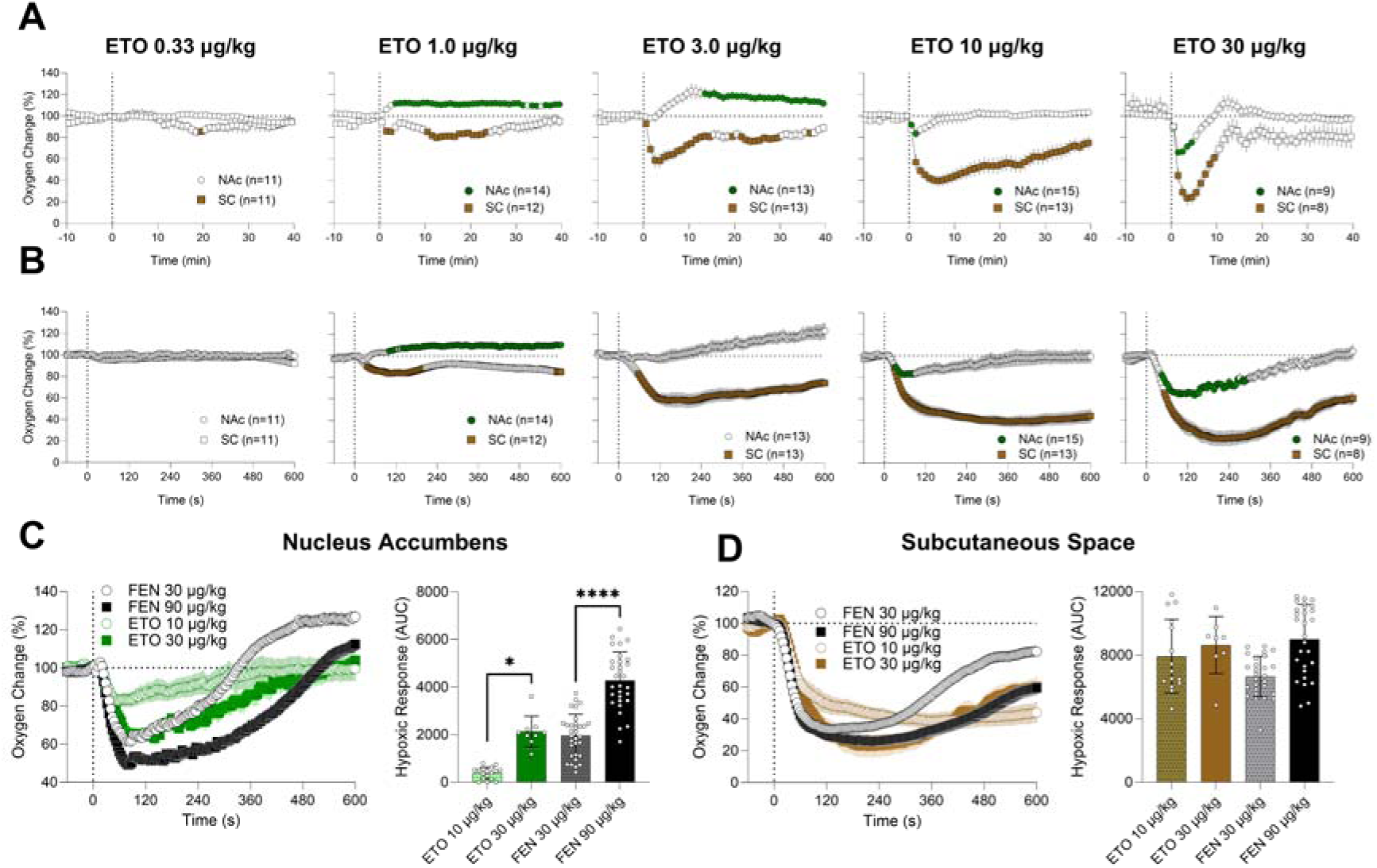
Mean (±SEM) changes in oxygen levels induced by iv etonitazene (ETO) in the NAc and subcutaneous space at different doses. A shows data analyzed with slow (1-min) time resolution. B shows data analyzed with rapid (4-s) time resolution. C show between-drug differences in brain oxygen levels analyzed with high temporal resolution (left graph) and changes in hypoxic effects of drug analyzed using area under the curve (right bar-graph; asterisks show differences with p<0.05, p<0.01, p<0.001, and p<0.0001). D show similar changes in the subcutaneous space. Results of statistical evaluations for A and B are shown in Supplementary materials (S2). Filled symbols show values significantly different from baseline (p<0.05). n = numbers of averaged tests.

Changes in oxygen levels induced by ETO have several important differences from responses induced by FEN (Figure 2C and D). To evaluate these differences, we compared the patterns and mean values of oxygen decreases (AUC) induced by ETO and FEN at two large doses (FEN 30 and 90 µg/kg; ETO 10 and 30 µg/kg). Brain hypoxia induced by ETO at a 10 µg/kg dose was relatively weak in magnitude, duration, and area under the curve. At a 30 µg/kg ETO dose, the decrease was much stronger and longer in duration. These changes were similar to those induced by FEN at the same dose, but weaker than changes induced by FEN at the largest dose. In contrast, oxygen decreases induced in the subcutaneous space by both drugs were relatively similar in their magnitude and in their area under the curve.

### 3.3. Brain and peripheral oxygen fluctuations induced by fluornitrazene (FNZ)

Figure 3 shows the effects of FNZ on oxygen levels in the NAc and SC space (see Supplementary table S3 for results of statistical evaluations). These effects were very similar to the effects of FEN at the similar dose. At the lowest effective, 1 µg/kg dose, FNZ induced a weak increase in brain oxygen levels, which became stronger at a 3 µg/kg dose. At this dose, we observed a weak but significant oxygen decrease in the subcutaneous space. At the two highest, 10 and 30 µg/kg doses, FNZ induced biphasic brain oxygen responses, with an initial decrease followed by rebound-like increase. The decrease was relatively weak (mean at nadir =74.96% and 57.39% of baseline for 10 and 30 µg/kg doses, respectively) and relatively short in duration (3-8 min). In the subcutaneous space FNZ induced a monophasic oxygen decrease, which was stronger than that in the brain (mean=54.77% and 48.78% for 10 and 30 µg/kg, respectively).

**Fig. 3.**
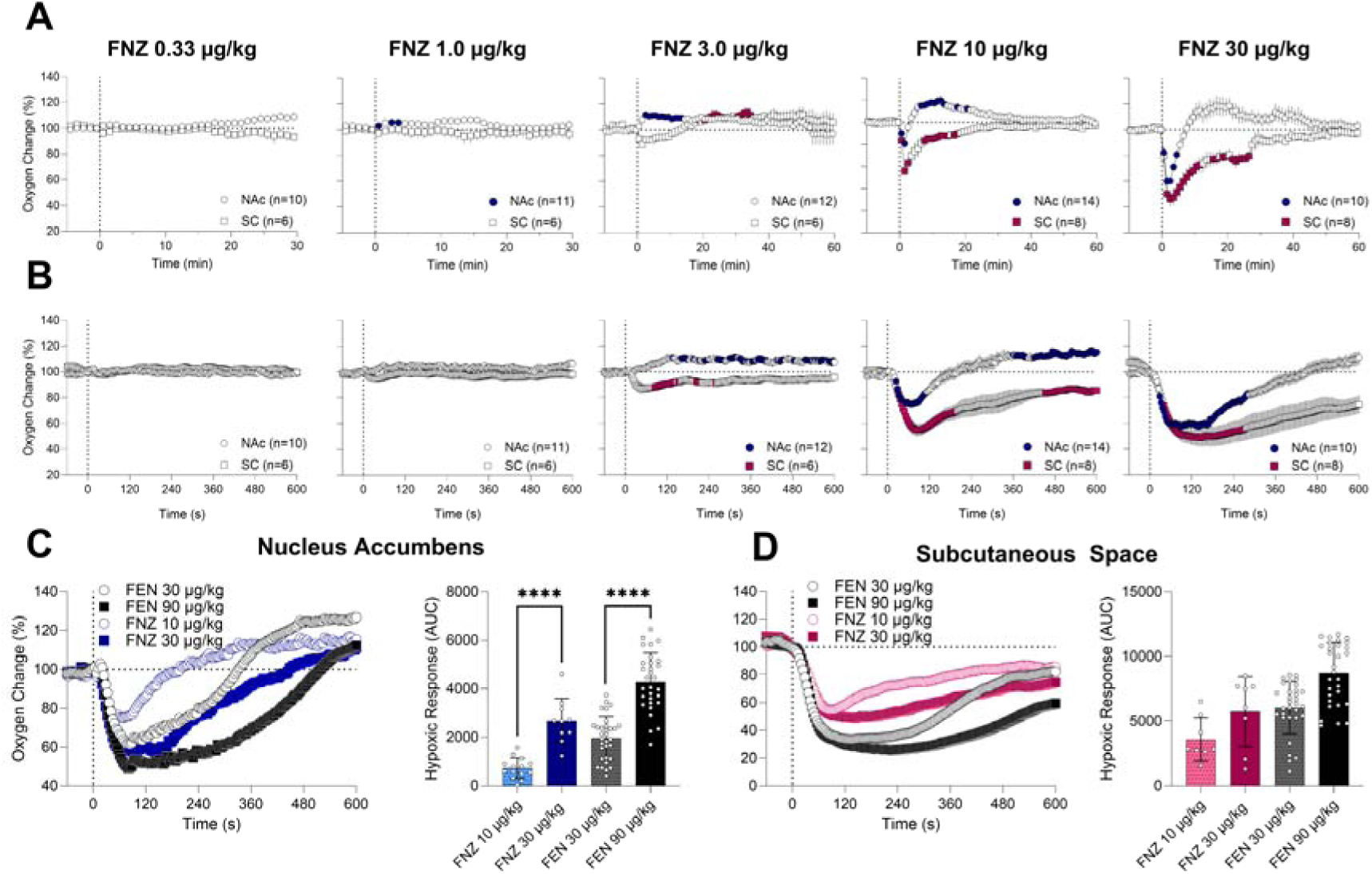
Mean (±SEM) changes in oxygen levels induced by iv FNZ in the NAc and subcutaneous space at different doses. A shows data analyzed with slow (1-min) time resolution. B shows data analyzed with rapid (4-s) time resolution. C show between-drug differences in brain oxygen levels analyzed with high temporal resolution (left graph) and changes in hypoxic effects of drug analyzed as the area under the curve (right graph) (asterisks show differences with p<0.05, p<0.01, p<0.001, and p<0.0001). D show similar changes in the subcutaneous space. Results of statistical evaluations for A and B are shown in Supplementary materials (S3). Filled symbols show values significantly different from baseline (p<0.05). n = numbers of averaged tests.

Brain oxygen decreases induced by FNZ showed great similarity with the effects of FEN (Figure 3C and 3D). While the area under the curve was significantly larger for higher doses of both drugs, this parameter was similar for both drugs at equal doses of 30 µg/kg. In the subcutaneous space, differences between the two drugs were minimal across doses.

### 3.4. Brain and peripheral oxygen fluctuations induced by N-desethyl-fluornitrazene (DFNZ)

DFNZ induced a quite different pattern of oxygen responses than FEN, ETO, and its parent compound FNZ (Figure 4A and B; see Supplementary table S4 for results of statistical evaluations). At the two low,10 and 30 µg/kg doses, DFNZ induced very weak and transient changes in oxygen: decreases in the subcutaneous space and biphasic changes in the NAc consisting of initial decreases followed by increases. At a higher, 100 µg/kg doses, subcutaneous decreases became stronger and more prolonged, and changes in the NAc were minimal and with a weak tonic increase. When DFNZ was injected at a very high, 300 µg/kg iv dose, we observed a weak but significant drop in NAc oxygen (mean=88.54% of baseline at nadir) followed by a much stronger and prolonged oxygen increase. At this dose, the decrease in subcutaneous space was clearly stronger and more prolonged (mean=75.08%) than in the NAc.

**Fig. 4.**
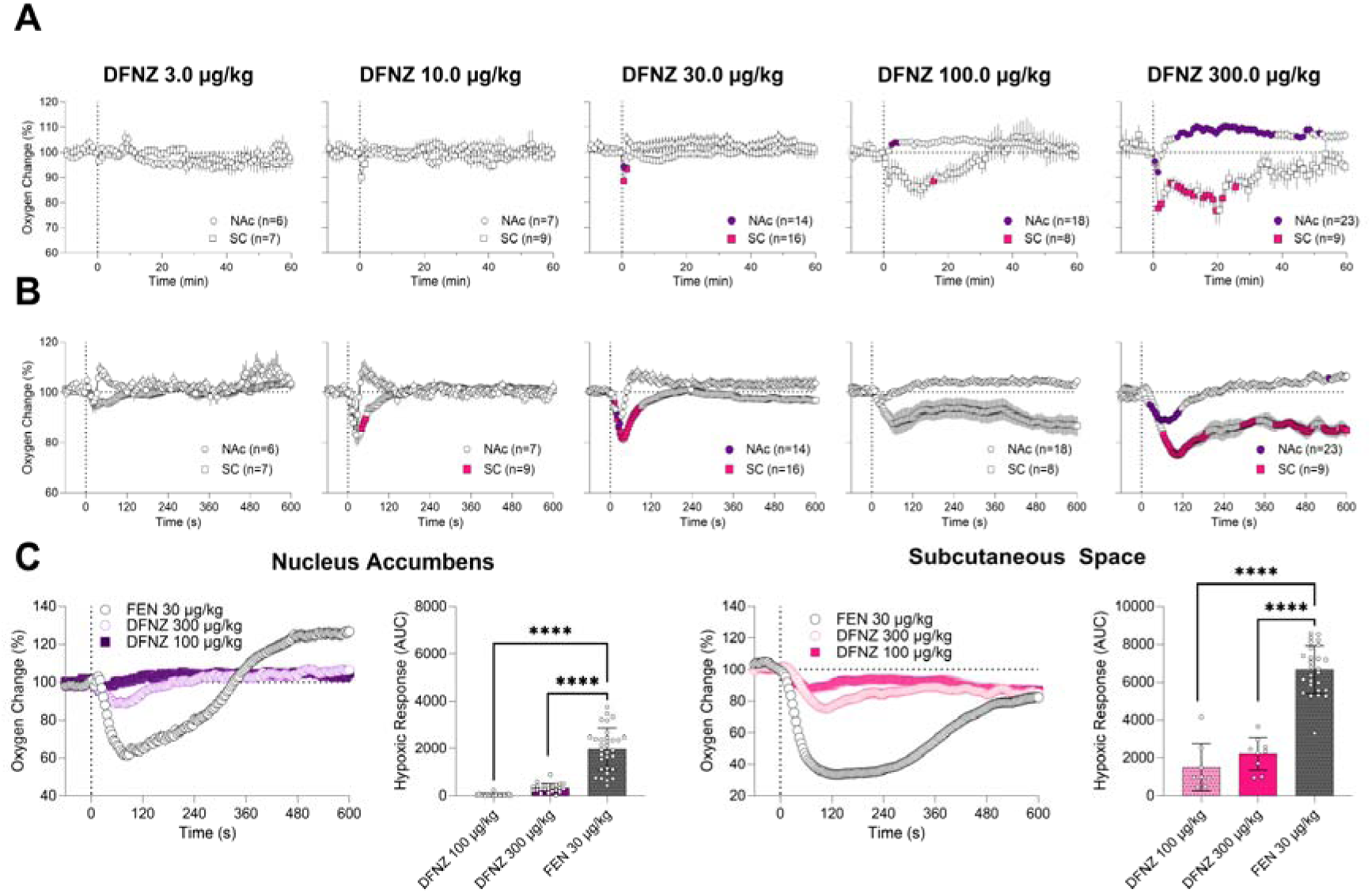
Mean (±SEM) changes in oxygen levels induced by iv DFNZ in the NAc and subcutaneous space at different doses. A shows data analyzed with slow (1-min) time resolution. B shows data analyzed with rapid (4-s) time resolution. C show between-drug differences in brain oxygen levels analyzed with high temporal resolution (left graph) and changes in hypoxic effects of drug analyzed as the area under the curve (right graph) (asterisks show differences with p<0.05, p<0.01, p<0.001, and p<0.0001). D show similar changes in the subcutaneous space. Results of statistical evaluations for A and B are shown in Supplementary materials (S4). Filled symbols show values significantly different from baseline (p<0.05). n = numbers of averaged tests.

Next, we compared the hypoxic effects of DFNZ at two high doses (100 and 300 µg/kg) with the effect of FEN at a moderate dose (30 µg/kg), which induced strong biphasic brain oxygen responses. As can be seen in Figure 4C, DFNZ has virtually no hypoxic effects on NAc oxygen levels at a 100 µg/kg dose and minimal changes at the highest dose (300 µg/kg). DFNZ at these two high doses also had minimal effects on oxygen levels in the subcutaneous space (Figure 4D).

To confirm the unusually weak effects of DFNZ, we examined brain oxygen responses at a very large, 1.0 mg/kg iv dose, which is about 30-fold higher than the maintenance self-administering dose (Gomez et al., 2026). At this dose, DFNZ induced profound sedation, but tail erection and increased muscular tone were not clearly observed. As can be seen in Figure 5A (see Supplementary table S5 for results of statistical evaluations), DFNZ at this large dose induced brain hypoxia, although it was relatively short-lived (∼6 min); this effect was similar in its latency and nadir to the effects of FNZ, its parent drug, at a very low dose (10 µg/kg).

**Fig. 5.**
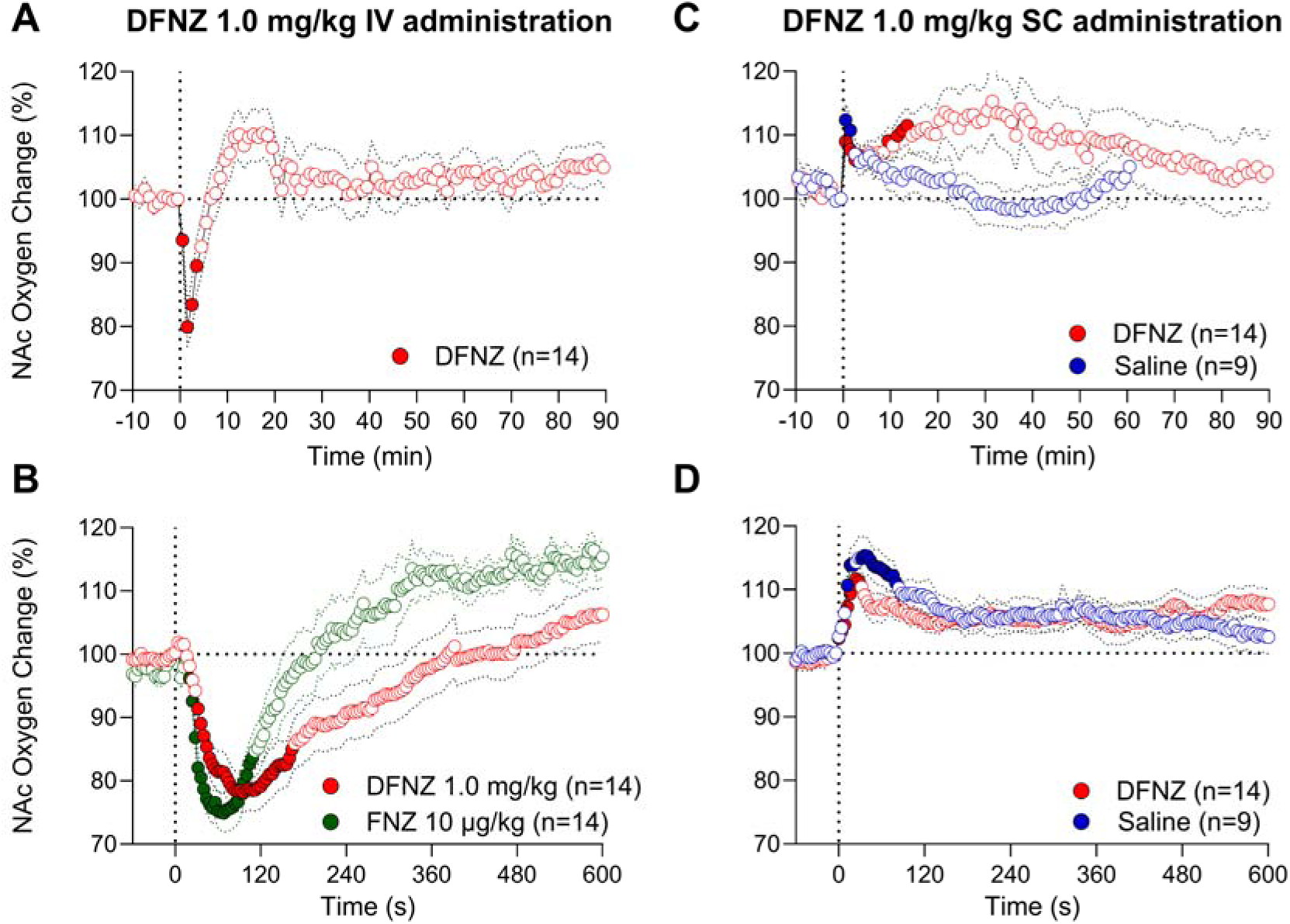
**A-B**. Mean (±SEM) changes in NAc oxygen responses induced by iv injection of DFNZ at a 1.0 mg/kg dose. Data analyzed with slow, 1-min (A) and rapid, 4-s (B) time resolution. **C-D**. Mean (±SEM) changes in oxygen responses induced by subcutaneous injection of saline (blue) and DFNZ at a higher 1.0 mg/kg dose (red). Data analyzed with slow, 1-min (C) and rapid, 4-s (D) time resolution. Filled symbols in all graphs show values significantly different from baseline, n is the number of averaged tests, and vertical hatched lines show the time of injection. Graph B shows brain oxygen response induced by FNZ at the dose 100-fold lower than that of DFNZ. See Supplementary table S5 for results of statistical evaluations.

As most studies assess a drug’s analgesic properties using a subcutaneous route of administration, we conducted additional experiments to examine how the effects of DFNZ depend on the route of administration. A high dose of 1.0 mg/kg was used to compare brain oxygen responses induced via subcutaneous and iv administration. In contrast to rapid oxygen increase after iv drug administration (Figure 5A, B), the effect of subcutaneous DFNZ delivery was biphasic, with a rapid short latency-induced increase following injection and subsequent weak tonic increase (Figure 5C, D). However, a very similar brain oxygen increase was seen after saline injection (both DFNZ and saline were delivered at a volume of 0.5 ml) suggesting the procedure of drug administration, but not the drug *per se*, is the cause of an initial, stress-related oxygen increase.

### 3.5. Area under the curve: comparisons for all drugs

Finally, to compare the hypoxic potency of each drug, we examined differences in oxygen decreases in the NAc and subcutaneous space using area under the curve (AUC) analysis (Figure 6). For all drugs, hypoxia was stronger in the subcutaneous space than in the NAc, and the effects of each drug strengthened with increases in drug dose. Three drugs, FEN, ETO and FNZ, showed similar high potency, inducing NAc hypoxia at a 10 µg/kg dose. Similar between-drug differences were seen with SC oxygen decreases. In this case, the effects of ETO were strongest, exceeding the effects of FEN and FNZ. Minimal potency to induce hypoxia was found with DFNZ, which produced the weakest oxygen decreases at 100 and 300 µg/kg doses. The AUC for oxygen decreases induced by DFNZ in the NAc at the largest dose (i.e., 300 µg/kg) (AUC 323) was 6-fold smaller than the AUC for FEN at a ten-fold lower FEN dose, 30 µg/kg dose (AUC 1,965), suggesting an at least 60-fold weaker potency of DFNZ in its ability to induce brain hypoxia compared to FEN. The ratio in hypoxic potency was even larger for DFNZ at a 100 µg/kg dose (∼150). However, the hypoxic effect of DFNZ in the subcutaneous space was larger than in the NAc (AUC: 2204 vs. 323) and was similar for both high DFNZ doses.

**Fig. 6.**
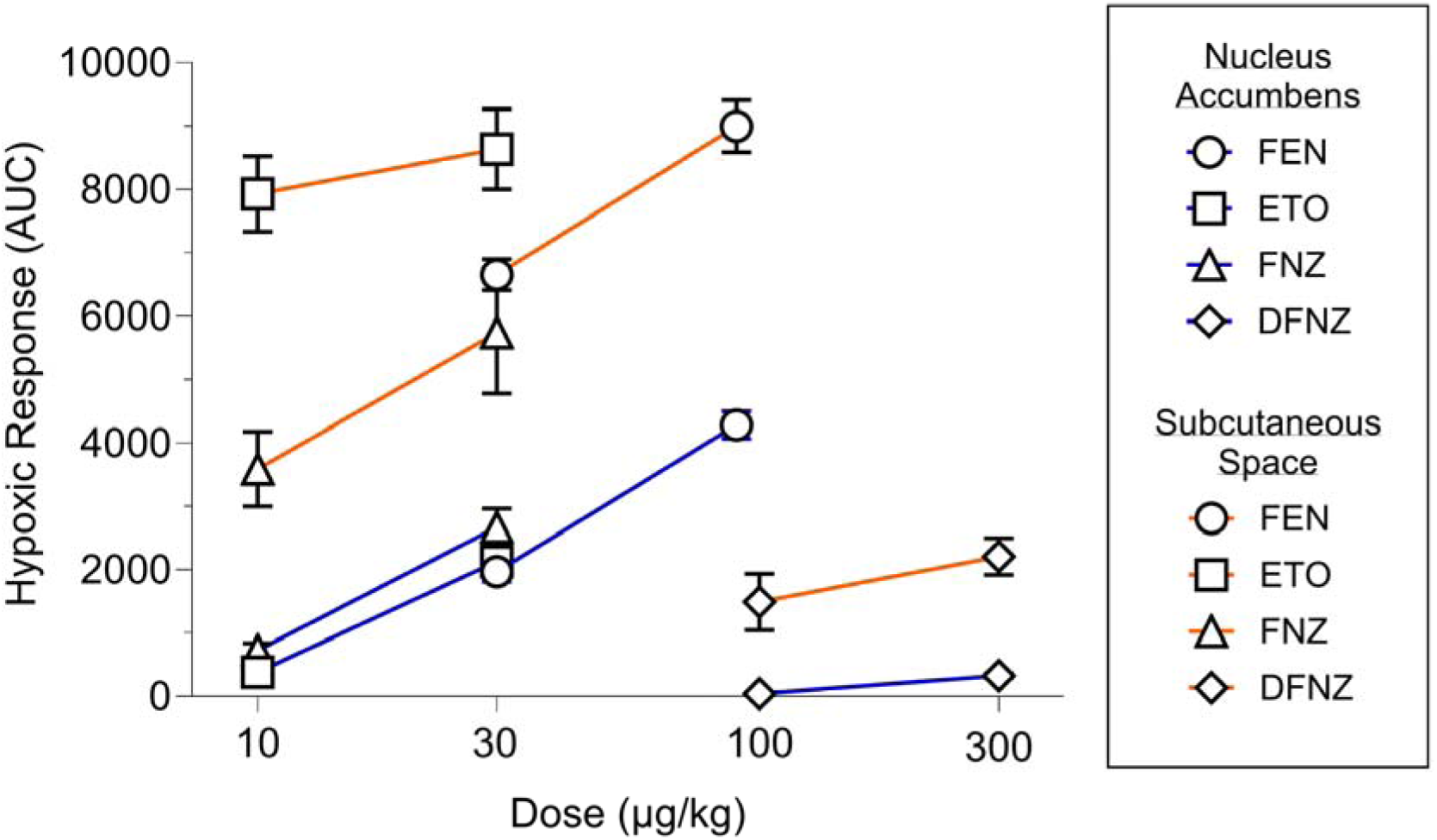
Mean (±SEM) changes in oxygen levels induced by iv injections of fentanyl, etonitazene, FNZ and DFNZ. Data analyzed as area under the curve for oxygen decreases and shown using a natural logarithmic scale. Orange lines show changes in the subcutaneous space and blue lines show changes in the NAc.

## 4. Discussion

### 4.1. Methodological considerations

The primary goal of this study was to examine hypoxic effects of classic and novel compounds of the nitazene family, a major factor determining the toxicity of opioids. Since dramatic changes in breathing are the primary cause of drug-induced hypoxia, most studies assessing drug-induced toxicity have focused on monitoring respiratory activity using plethysmography. While plethysmography provides essential clues to the hypoxic effects of opioids, respiratory activity is a peripheral parameter and does not provide direct insights into the status of brain oxygen dynamics, which are more tightly coupled to toxicity than respiration. Electrochemical technology addresses this problem, by directly assessing drug-induced oxygen responses in the brain and peripheral locations simultaneously and with greater, second-scale temporal resolution in freely moving rats.

### 4.2. Common features and differences in oxygen fluctuations induced by nitazenes

All nitazenes used in this study induced brain hypoxia, but this effect was dose-dependent, and the pattern had both common and distinguishing features from the hypoxia induced by FEN. Similar to our previous studies with heroin and FEN (Thomas et al., 2021; Curay et al., 2023; Choi et al., 2024), all opioid drugs in this study induced moderate but significant brain oxygen *increases* when used at very low, pre-hypoxic doses. Although this finding can be viewed as paradoxical, opioids injected in very small doses interact with opioid receptors, likely mimicking the effects induced by endogenous opioid peptides, which are released by different arousing and stress-inducing stimuli under normal conditions (Drolet et al., 2001; Higginbotham et al., 2022). Therefore, opioids, depending on the dose, may induce opposite brain oxygen responses. Considering this factor, the minimal effective doses of ETO, FNZ, and FEN, which induced brain hyperoxic responses were between 1 and 3 µg/kg, clearly lower than doses inducing hypoxic responses (between 3 and 10 µg/kg), but within the range for maintaining self-administration behavior (FEN, Malone et al., 2021; ETO, Carroll and Boe, 1984; Gomez, and Meisch, 2004; FNZ, Gomez et al., 2026).

Hypoxic responses were also observed in the periphery, and these effects were always stronger and more prolonged than the responses seen in the brain. The oxygen dynamics induced by all opioids in the periphery had two important differences from brain oxygen responses. While oxygen responses in the brain were biphasic, with rapid, strong, and relatively brief decreases followed by more prolonged post-hypoxic oxygen increases (i.e., hyperoxia), oxygen responses in the periphery were monophasic, with rapid decreases and slower return to baseline. Since respiratory depression results in intra-cerebral accumulation of CO_2_, a powerful vasodilator (Schmidt and Kety, 1947; Kontos, 1981, Battisti-Charbonney et al., 2011), cerebral vasodilation and increased global cerebral blood flow may contribute to the post-hypoxic oxygen increases as an adaptive brain mechanism opposing hypoxia. This effect appears to be unique for the brain, as it was absent in the periphery. In contrast, oxygen decreases in the periphery result from the summation of two factors: respiratory depression and peripheral vasoconstriction, which decreases oxygen inflow into peripheral tissues (Solis et al., 2018; Thomas et al., 2021). Therefore, the stronger and protracted peripheral hypoxia produced by all drugs may reflect the capacity of the drugs to not just inhibit respiration but also to produce vasoconstriction.

### 4.3. ETO as a highly potent member of the nitazene family

Early studies conducted in mice, dogs, and monkeys revealed that ETO is highly toxic and potent in inducing respiratory depression (Butelman et al., 1993; Carregal and Williams, 1967; Elmer et al., 1995). Consistent with these data, we found that iv delivery of ETO induced brain oxygen decreases, which were evident at a low,10 µg/kg dose. Notably, while the brain hypoxic effect of ETO assessed by AUC was similar to the effect of FEN (see Figure 6), the duration of brain hypoxia induced by this drug was clearly more prolonged than that of FEN. Also, a prominent post-hypoxic hyperoxia typical of FEN, FNZ and DFNZ, was virtually absent in the effects of ETO.

It is known that the damaging effects of hypoxia on brain cells and neural functions exponentially increase with prolongation of hypoxia (Lee et al., 2000; Bailey, 2019; Terraneo and Samaja, 2017). Brain adaptive mechanisms, within some limits, can oppose strong but transient hypoxia, however brain damage progresses when hypoxia becomes prolonged. Thus, more prolonged hypoxia and the absence of post-hypoxic hyperoxia could be a factor determining the high toxicity of ETO. While not objectively monitored, ETO induced stronger abnormal behavioral effects than FEN, including extreme tail erection, strong muscle rigidity, deep sedation, gasping, and convulsions seen in some rats. Due to these clear adverse effects, the largest dose of ETO in our tests was limited to 30 µg/kg, although FEN produced similar or weaker behavioral effects at a 90 µg/kg dose.

The threshold doses of ETO to induce brain hypoxia in this study (3-10 µg/kg) are within the range of analgesic doses as evaluated by tail-flick and hot plate tests in mice (Ujváry et al., 2021; Gatch et al., 2025). While we were unable to find data obtained in rats with iv drug delivery, similar thresholds of effective analgesic doses of ETO (ED_50_=9 µg/kg) were reported with subcutaneous administration in mice (Ujváry et al., 2021; Gatch et al., 2025). In these studies, the effect appeared at 10 µg/kg and became stronger at 30 µg/kg (ED_50_=22 µg/kg). However, data obtained in rats using the hot plate technique revealed stronger analgesic potency of subcutaneous ETO that exceeds the values for FEN (ED_50_=1.7 vs. 21 µg/kg) (Vandeputte et al., 2023). In this study, a ten-fold higher potency of ETO vs. FEN was also observed when assessing catalepsy, another prominent effect of opioids.

Although ETO is considered highly toxic, with its toxicity exceeding that of FEN, experimental data are very limited and therefore this notion is controversial. For example, ED_50_ values obtained in mice with subcutaneous or intraperitoneal administration were higher (ED_50_=1 mg/kg; Ujváry et al., 2021) compared to values inducing robust adverse effects following iv administration in rats. The reasons for these differences remain unclear, and this issue requires further experimental testing. We were unable to find any data on toxicity of ETO in rats following iv administration, but studies have shown that rats can reliably, and safely, self-administer ETO at doses up to 80 µg/kg (Carrol et al., 1981). Nevertheless, since ETO induced robust behavioral effects at a 30 µg/kg iv dose, we can estimate that doses exceeding this value could be potentially toxic.

### 4.4. FNZ and DFNZ: Fluorated nitazane analogs related to ETO

Fluornitrazene (FNZ) is a fluorinated derivative of ETO with high affinity at µ-opioid receptors (K_i_=1.36 nM) and superagonist efficacy (Gomez et al., 2026). Our findings showed that FNZ produced similar effects on oxygen homeostasis as FEN. Like FEN, brain oxygen levels increased at lower iv doses (1-3 µg/kg) and produced biphasic responses at a moderate and higher dose (10 and 30 µg/kg). Specifically, oxygen decreases appeared at a 3 µg/kg dose, became stronger and more prolonged at 10 µg/kg, and further increased at a 30 µg/kg dose. The hypoxic threshold of FNZ in the brain (∼6 µg/kg) is similar to its optimal self-administered dose (10 µg/kg) and analgesic threshold (ED_50_=5 µg/kg) as assessed using iv self-administration and the hot plate technique in rats with subcutaneous delivery respectively (Gomez et al., 2026).

DFNZ is a highly selective super-agonist of the μ-opioid receptor, with an affinity (K_i_) of 1.0 nM, which is higher than affinities of both ETO (Ki= ∼8.11 nM) and FEN (Ki=6.35 nM) (Gomez et al., 2026). It exhibits brain permeability but reduced capacity to cross the blood–brain barrier which leads to restricted brain u-opioid receptor occupancy.

Our results showed that DFNZ has quite unusual effects on oxygen levels, differing from the effects of FEN, ETO, and even its parent, FNZ. Within a wide range of doses, DFNZ had no evident effects on brain oxygen levels except for a weak but significant decrease at a large, 100 µg/kg dose. Oxygen changes in the periphery were also very weak and brief, producing small dose-dependent decreases. Clear effects of DFNZ in the brain were seen only at very large, 300-1000 µg/kg doses, 10-30-fold higher than the maintenance self-administering dose (Gomez et al., 2026). At these doses, DFNZ had also a clear effect in the periphery, inducing a tonic oxygen decrease. Hypoxic effects of this drug, as assessed by decreases in mean response magnitude and AUC, were also quite weak, within the range of peripheral oxygen fluctuations consistently seen in awake rats during day-time recordings. In contrast to FNZ, the doses of DFNZ to induce brain hypoxia were much larger with a ∼60-fold between-drug ratio. Interestingly, the very same DFNZ-FNZ ratio (∼60-fold) was found when assessing analgesia using the hot plate after subcutaneous injection (ED_50_=300 µg/kg vs. 5 µg/kg in FNZ).

Since most analgesic tests are conducted using subcutaneous drug delivery, we compared the effects of DFNZ at a very large, 1000 µg/kg dose induced by sc injections. While iv-delivered DFNZ induced weak and transient brain oxygen decreases, such decreases after sc administration were absent and brain oxygen levels showed two-component *increases*, with a phasic change after the injection and subsequent much weaker and more tonic increase. As shown for FEN, these differences in oxygen dynamics obviously result from differences in drug pharmacokinetics, with very rapid (1-2 min) peaks in drug levels after iv injection and definite onset latencies, much lower peaks, and longer duration after subcutaneous delivery (Ohtsuka et al., 2007; Schleimer et al., 1978; MacKenzie et al., 2016). However, we found that saline injection induces a similar post-injection phasic oxygen increase, suggesting that the stressful procedure of drug administration, but not necessarily the drug itself, is a possible cause of this change. However, the weak and delayed oxygen increase could be drug-induced, but it is unclear whether this small change is real increase or spontaneous oxygen fluctuation. This state-dependency appears to be common for other drugs as the effects of FEN were also strongly dependent on administration route. While iv-delivered FEN induced clear hypoxic responses at a 30 µg/kg dose, the hypoxic effect of this drug delivered as intraperitoneal injection at much higher, 300 µg/kg dose was comparable, even slightly lower in magnitude and slightly longer in duration (Limiac et al., 2025). The latter dose is clearly within the range of possible lethality in rats (Lewis et al., 2026; Boom et al., 2012; Strauss et al., 2024). Therefore, the route of drug administration, could be an important contributor to high variability in quantitative parameters of hypoxic effects and general toxicity of opioid drugs, which also have species specificity.

The high temporal resolution of our oxygen recordings revealed that DFNZ at relatively low doses (10-30 µg/kg) also had rapid and brief effects on brain oxygen levels, which had ultra-short onset latencies. While the underlying mechanism remains unclear, this pattern may suggest direct actions of DFNZ on opioid receptors on afferents of sensory nerves densely innervating blood vessels in peripheral tissues (Stein, 2013, 2016; Smith 2008; Coggeshall et al., 1997). This transient effect was evident only with low-dose DFNZ injections and not seen with higher drug doses due to opposing weak hypoxic effect of this drug.

In conclusion, we show that several nitazene analogs differ in their capacity to produce hypoxia in the brain and in peripheral tissues. Specifically, the novel nitazene DFNZ showed a divergent hypoxia profile when compared to ETO, the prototypical compound of this class, FNZ, its parent molecule, and FEN. Thus, by using simultaneous recordings from the NAc and subcutaneous space, we showed that for each tested drug, hypoxia was stronger and more prolonged in the peripheral tissues than in the brain due to brain-specific adaptive mechanisms. Finally, we showed that minor changes to the molecular structure of nitazene compounds result in profound differences in their hypoxic effects. While FNZ resembled the hypoxic effects of ETO and FEN, DFNZ showed a remarkably weak hypoxic profile, supporting the notion that some nitazenes may have less potential for toxicity than others.

## Author contributions: CRediT

EAK: Conceptualization, Data curation, formal analysis, funding acquisition, investigation, methodology, Writing-original draft, Writing-review and editing; AA and NN: surgeries, performance of experiments, data analyses, histological work, data curation; MM and JG: conceptualization, writing-original draft, Writing-review and editing the manuscript. AS, KCR: synthesis of novel substances (FNZ, DFNZ). All authors were involved in manuscript’s writing and editing and all authors approved the final version.

## Funding

This research was supported by the Intramural Research Program of the National Institutes of Health (EAK). The contributions of the NIH authors are considered Works of the United States Government. The findings and conclusions presented in this paper are those of the authors and do not necessarily reflect the views of the NIH or the U.S. Department of Health and Human Services. Grant funding: 1ZIADA000566-03 (for Eugene A. Kiyatkin) and ZIADA000069 (for Michael Michaelides), and ZIADA000527 (for Kenner C. Rice).

## Declaration of competing interest

The authors declare that they have no known competing financial interests or personal relationships that could have appeared to influence the work reported in this paper.

## Data Availability Statement

Raw data and the results of their primary analyses are available on request from Eugene A. Kiyatkin (NIDA-IRP, NIH)

## Ethical Approval

The studies described in this paper were approved by the Animal Care and Use Committee of the Intramural Research Program of NIDA, NIH.

## Acknowledgements

We want to acknowledge the assistance of three post-bac fellows (Shinbe Choi, Michael R. Noya, Feonil G. Limiac) with FEN data collection and analysis.

## Notes

### Competing Interest Statement

The authors have declared no competing interest.

