## Supplementary Material for "Distinct changes in brain and peripheral oxygen levels induced by synthetic opioids of the nitazene family"

|  | **Fig 1A. NAc** | **Fig 1A. SC Space** | **Fig 1B. NAc** | **Fig 1B. SC Space** |
| --- | --- | --- | --- | --- |
| FEN 1.0 ug/kg | F_24,744_=2.60* | F_21,651_=1.98^ns^ | F_24,3624_=2.05^ns^ | F_21,3171_=2.13^ns^ |
| FEN 3.0 ug/kg | F_26,1066_=7.83**** | F_24,984_=6.35**** | F_26,3926_=1.36^ns^ | F_24,3624_=3.58* |
| FEN 10 ug/kg | F_27,1647_=77.74**** | F_24,1440_=22.13**** | F_27,4077_=13.70**** | F_24,3624_=17.56**** |
| FEN 30 ug/kg | F_31,2821_=30.39**** | F_26,2366_=46.84**** | F_31,4681_=109.60**** | F_26,3926_=72.22**** |
| FEN 90 ug/kg | F_29,2639_=67.80**** | F_27,2457_=55.85**** | F_29,4379_=95.48**** | F_27,4077_=52.89**** |

**Table S1**. (Figure 1A and 1B). Results of statistical evaluation of changes in oxygen levels in the NAc and SC space induced by FEN * p<0.05; **, p<0.01; ***, p<0.001, **** p<0.0001.

|  | **Fig 2A. NAc** | **Fig 2A. SC Space** | **Fig 2B. NAc** | **Fig 2B. SC Space** |
| --- | --- | --- | --- | --- |
| ETO 0.33 ug/kg | F_10,410_=3.01* | F_10,410_=2.82* | F_10,1510_=0.78^ns^ | F_10,1510_=1.01^ns^ |
| ETO 1.0 ug/kg | F_13,533_=2.51^ns^ | F_11,451_=2.04^ns^ | F_13,1963_=8.11*** | F_11,1661_=3.12^ns^ |
| ETO 3.0 ug/kg | F_12,732_=5.59* | F_12,732_=8.67**** | F_12,1812_=11.34**** | F_12,1812_=13.66**** |
| ETO 10 ug/kg | F_14,1274_=1.90^ns^ | F_14,1274_=9.98**** | F_14,2114_=3.02^ns^ | F_14,2114_=16.58*** |
| ETO 30 ug/kg | F_8,728_=10.00**** | F_7,637_=8.06** | F_8,1208_=15.37**** | F_7,1057_=16.61**** |

**Table S2**. (Figure 2A and 2B). Results of statistical evaluation of changes in oxygen levels in the NAc and SC space induced by ETO * p<0.05; **, p<0.01; ***, p<0.001, **** p<0.0001.

|  | **Fig 3A. NAc** | **Fig 3A. SC Space** | **Fig 3B. NAc** | **Fig 3B. SC Space** |
| --- | --- | --- | --- | --- |
| FNZ 0.33 ug/kg | F_9,279_=5.34** | F_5,155_=1.13^ns^ | F_9,1359_=1.21^ns^ | F_5,755_=1.23^ns^ |
| FNZ 1.0 ug/kg | F_10,310_=1.98^ns^ | F_5,155_=0.62^ns^ | F_10,1510_=1.10^ns^ | F_5,755_=0.68^ns^ |
| FNZ 3.0 ug/kg | F_11,671_=1.94^ns^ | F_5,305_=4.45* | F_11,1661_=3.86** | F_5,755_=2.76^ns^ |
| FNZ 10 ug/kg | F_13,793_=7.65**** | F_7,427_=12.20** | F_13,1063_=27.06**** | F_7,1057_=8.78** |
| FNZ 30 ug/kg | F_9,549_=19.36**** | F_7,427_=20.00**** | F_9,1359_=44.34**** | F_7,1057_=7.25* |

**Table S3**. (Figure 3A and 3B). Results of statistical evaluation of changes in oxygen levels in the NAc and SC space induced by FNZ * p<0.05; **, p<0.01; ***, p<0.001, **** p<0.0001.

|  | **Fig 4A. NAc** | **Fig 4A. SC Space** | **Fig 4B. NAc** | **Fig 4B. SC Space** |
| --- | --- | --- | --- | --- |
| DFNZ 3.0 ug/kg | F_5,305_=2.25^ns^ | F_6,366_=1.34^ns^ | F_5,755_=1.68^ns^ | F_6,906_=2.13^ns^ |
| DFNZ 10 ug/kg | F_6,366_=1.06^ns^ | F_8,488_=1.88^ns^ | F_6,906_=3.68* | F_8,1208_=9.45*** |
| DFNZ 30 ug/kg | F_13,793_=1.84^ns^ | F_15,915_=2.45^ns^ | F_13,1963_=5.93**** | F_15,2265_=28.29**** |
| DFNZ 100 ug/kg | F_17,1073_=1.99^ns^ | F_7,427_=4.76* | F_17,2567_=6.42**** | F_7,1057_=2.66^ns^ |
| DFNZ 300 ug/kg | F_22,14342_=7.90**** | F_8,488_=4.73** | F_22,3322_=18.76**** | F_8,1208_=6.27**** |

**Table S4**. (Figure 4A and 4B). Results of statistical evaluation of changes in oxygen levels in the NAc and SC space induced by DFNZ * p<0.05; **, p<0.01; ***, p<0.001, **** p<0.0001.

| **6A.** | **NAc** |
| --- | --- |
| DFNZ 1.0 mg/kg IV (1-min resolution) | F_18,648_=10.44**** |
| DFNZ 1 mg/kg (4-s resolution) | F_18,1728_=10.63**** |
| FNZ 1 mg/kg (4-s resolution) | F_13,1063_=27.06*** |
| **6B.** |  |
| DFNZ 1.0 mg/kg SC (1-min resolution) | F_13,793_=1.56^ns^ |
| Saline SC (1-min resolution) | F_8,248_=3.05^ns^ |
| DFNZ 1.0 mg/kg SC (4-s resolution) | F_13,1963_=1.31^ns^ |
| Saline SC (4-s resolution) | F_8,1208_=4.77** |

**Table S5**. (Figure 5A and 5B). Results of statistical evaluation of changes in oxygen levels in the NAc induced by i.v. and subcutaneous injection of 1 mg/kg DFNZ (with comparisons to saline and 10 ug/kg FNZ) * p<0.05; **, p<0.01; ***, p<0.001, **** p<0.0001.
